# Deciphering octoploid strawberry evolution with serial LTR similarity matrices for subgenome partition

**DOI:** 10.1101/2024.07.31.606053

**Authors:** Haomin Lyu, Shujun Ou, Won Cheol Yim, Qingyi Yu

## Abstract

Polyploidization has been recognized as a major force in plant evolution. With the continuous progress in sequencing technologies and genome assembly algorithms, high-quality chromosome-level assemblies of polyploid genomes have become increasingly attainable. However, accurately delineating these assemblies into subgenomes remains a challenging task, especially in case where known diploid ancestors are absent. In this study, we introduce a novel approach that leverages long terminal repeat retrotransposons (LTR-RTs) coupled with the Serial Similarity Matrix (SSM) method to assign genome assemblies to subgenomes, particularly beneficial for those without known diploid progenitor genomes. The SSM method helps identify subgenome-specific LTRs and facilitates the inference of the timing of allopolyploidization events. We validated the efficacy of the SSM approach using well-studied allopolyploidy genomes, *Eragrostis tef* and *Gossypium hirsutum*, alongside an artificially created allotetraploid genome by merging two closely related diploid species, *Glycine max* and *G. soja*. Our results demonstrated the robustness of the method and its effectiveness in assigning chromosomes to subgenomes. We then applied the SSM method to the octoploid strawberry genome. Our analysis revealed three allopolyploidization events in the evolutionary trajectory of the octoploid strawberry genome, shedding light on the evolutionary process of the origin of the octoploid strawberry genome and enhancing our understanding of allopolyploidization in this complex species.

## Introduction

Polyploidization, or whole genome duplication (WGD), has been recognized as one of the major factors driving plant evolution (Ren et al. 2018; Soltis et al. 2014; Van de Peer et al. 2017). Nearly all seed plants have undergone at least one ancient WGD event (Tang et al. 2008; Jaillon et al. 2007; Bowers et al. 2003; Vanneste et al. 2014). Polyploidy is mainly classified into autopolyploidy and allopolyploidy based on the origin of chromosome set duplications (Spoelhof et al. 2017; Tayalé and Parisod 2013). Autopolyploids arise from within-species WGD, while allopolyploids result from interspecific hybridization followed by WGD. Approximately 24% of vascular plant species are polyploid, with 13% being autopolyploid and 11% allopolyploid (Wood et al. 2009; Barker et al. 2016). Allopolyploidy is notably prevalent in the Poaceae family, constituting 70 to 80% of all grasses (Kellogg and Kubitzki 2015; Stebbins 1985).

Allopolyploidization, combining diverse genetic compositions, plays an important role in plant speciation and evolution. Allopolyploid plants often exhibit broader resilience to harsh environments compared to their diploid counterparts, enabling survival in disasters and harsh climates, and gradual replacement of ecological niches, and even colonization of new habitats (Session et al. 2016; Sehrish et al. 2014; Yoo et al. 2013; Cox et al. 2014; Van de Peer et al. 2017, 2009, 2021).

Advancements in sequencing technologies and genome assembly algorithms now allow high-quality, chromosome-level assemblies of polyploid genomes. Genome sequencing of allopolyploid plants, such as tobacco (Sierro et al. 2014), teff (VanBuren et al., 2020), cotton (Hu et al., 2019), mustard (Yang et al. 2016; Yim et al., 2022), wheat (Appels et al. 2018), sugarcane (Zhang et al. 2018), strawberry (Edger et al. 2019), and coffee (Scalabrin et al. 2024, 2020), has significantly enhanced our understanding of allopolyploid evolution. Understanding structural and functional divergence between subgenomes is one of the most essential foundations for deciphering evolutionary innovation and ecological success of allopolyploids.

Identifying subgenomes is therefore critical for understanding evolutionary histories of allopolyploids. The commonly used method to identify subgenomes relies on the availability of genomic information from progenitor species. For an allotetraploid genome, at least one progenitor’s genome information is required to distinguish the two subgenomes. However, for many allopolyploid plants, the progenitor diploids are either unknown or extinct, making it difficult to assign sequence assemblies to subgenomes and trace their parental origins. Hence, there is a need to develop reliable bioinformatics pipelines for assigning chromosome-level assemblies into subgenomes in the absence of progenitor diploid genome sequences. Such tools are crucial for advancing our understanding of the role of polyploidization in plant evolution and speciation.

The cultivated strawberry (*Fragaria × ananassa*) is a recent allooctoploid (2n = 8x = 56) species. It is believed to have originated from a spontaneous hybridization between two octoploid progenitor species, *Fragaria chiloensis* and *F. virginiana* (Darrow 1966). However, cytological and phylogenetic studies yielded conflicting hypotheses regarding its diploid progenitors (Sargent et al. 2016; Senanayake and Bringhurst 1967; Tennessen et al. 2014; Yang and Davis 2017; Bringhurst 1990; Kamneva et al. 2017; DiMeglio et al. 2014; Rousseau-Gueutin et al. 2009; Fedorova 1946). Recent attempts identified four different diploid progenitor species for each subgenome of the cultivated strawberry, *Fragaria vesca* ssp. *bracteata*, *F. iinumae*, *F. viridis*, and *F. nipponica* using a tree-searching algorithm, PhyDS (Edger et al. 2019). However, the findings have been met with challenges and debate (Liston et al. 2020; Feng et al. 2020), highlighting the complexity of unraveling the evolutionary history of allopolyploid species. Liston et al. (Liston et al. 2020) revisited the analysis of the four subgenomes and found that the PhyDS method only supports the two extant diploid progenitors, *F. vesca* and *F. iinumae*, and no evidence supports *F. nipponica* and *F. viridis* ancestry or the hexaploidy species *F. moschata* as an evolutionary intermediate ancestor to the octoploid strawberry. Similarly, another study utilized an alignment-based approach and proposed *F. vesca* and *F. iinumae* as the diploid progenitors of *F.* x *ananassa*, while three other diploid species, *F. viridis*, *F. nilgerrensis*, and *F. nubicola*, were not considered parental species of *F. × ananassa* (Feng et al. 2020).

Transposable elements (TEs) are dynamic components of plant genomes, reflecting high diversity of lineages and distribution (Martelossi et al. 2023). TE dynamics are influenced by host regulations and responses to environmental stressors, potentially reflecting the evolutionary adaptation of their host organisms (Stapley et al., 2015; Lanciano and Mirouze 2018; Lisch 2013). Speciation events often coincide with the expansion of new TE families (Belyayev 2014; Jurka et al. 2011). Long terminal repeat retrotransposons (LTR-RTs) are the most abundant group of TEs in plant genomes, offering invaluable insights into the evolutionary history of plant species. A recent study investigating the allopolyploid grass *Leptochloa chinensis*, which lacks known parental species, suggests that transposable elements can be used to distinguish subgenomes (Wang et al. 2022). Session and Rokhsar further discussed the theoretical and empirical feasibility of leveraging subgenome-specific transposons for this purpose (Session and Rokhsar 2023). Moreover, subgenome-specific *k*-mers have been employed to assign chromosomes to subgenomes (Jia et al. 2022).

In this study, we introduce a bioinformatics pipeline for precise subgenome partition in allopolyploid genomes, particularly those lacking information on diploid progenitors. Our method utilizes the similarities of long terminal repeats (LTRs) to construct a Serial of Similarity Matrix (SSM), facilitating the effective clustering of chromosomes within the allopolyploid genome. Furthermore, by pinpointing subgenome-specific LTRs, the SSM enables inference regarding the timing of allopolyploidization events. We validated this approach using well-studied allopolyploidy genomes, tested it on an artificially constructed allotetraploid genome, and applied it to distinguish the subgenomes of the octoploid strawberry (*F. × ananassa* Duchesne) genome. Our findings elucidate the intricate evolutionary processes underlying allopolyploidization in this agriculturally important crop species.

## Results

### Development and validation of SSM method using well-studied allopolyploid genomes

Deciphering subgenome structure in allopolyploids lacking diploid progenitor information presents a significant challenge. However, LTR-RTs offer immense potential for addressing this issue due to their unique characteristics. Firstly, LTR-RTs are ubiquitous, comprising a substantial portion of plant genomes (Hawkins et al. 2009). Secondly, these elements possess the ability to replicate and relocate within the genome, potentially dispersing throughout the genome (Bourque et al. 2018). Thirdly, even after inactivation, LTR remnants persist, serving as valuable signatures of past genomic events (Lyu et al. 2018). Furthermore, LTR relics generally experience neutral or negative selection (Pereira 2004). Considering the evolutionary timeline of allopolyploids, LTR-RTs that emerged between the divergence of progenitor species and the allopolyploidization event are expected to be lineage-specific, offering distinct features for subgenome identification (Fig. 1A). We designate this stage “Phase 2”. To identify “Phase 2” for chromosome clustering and subgenome partitioning, we constructed a similarity matrix of LTRs at different cutoffs, resulting in a Serial of Similarity Matrix (SSM) (Fig. 1A and 1B).

**Figure 1.**
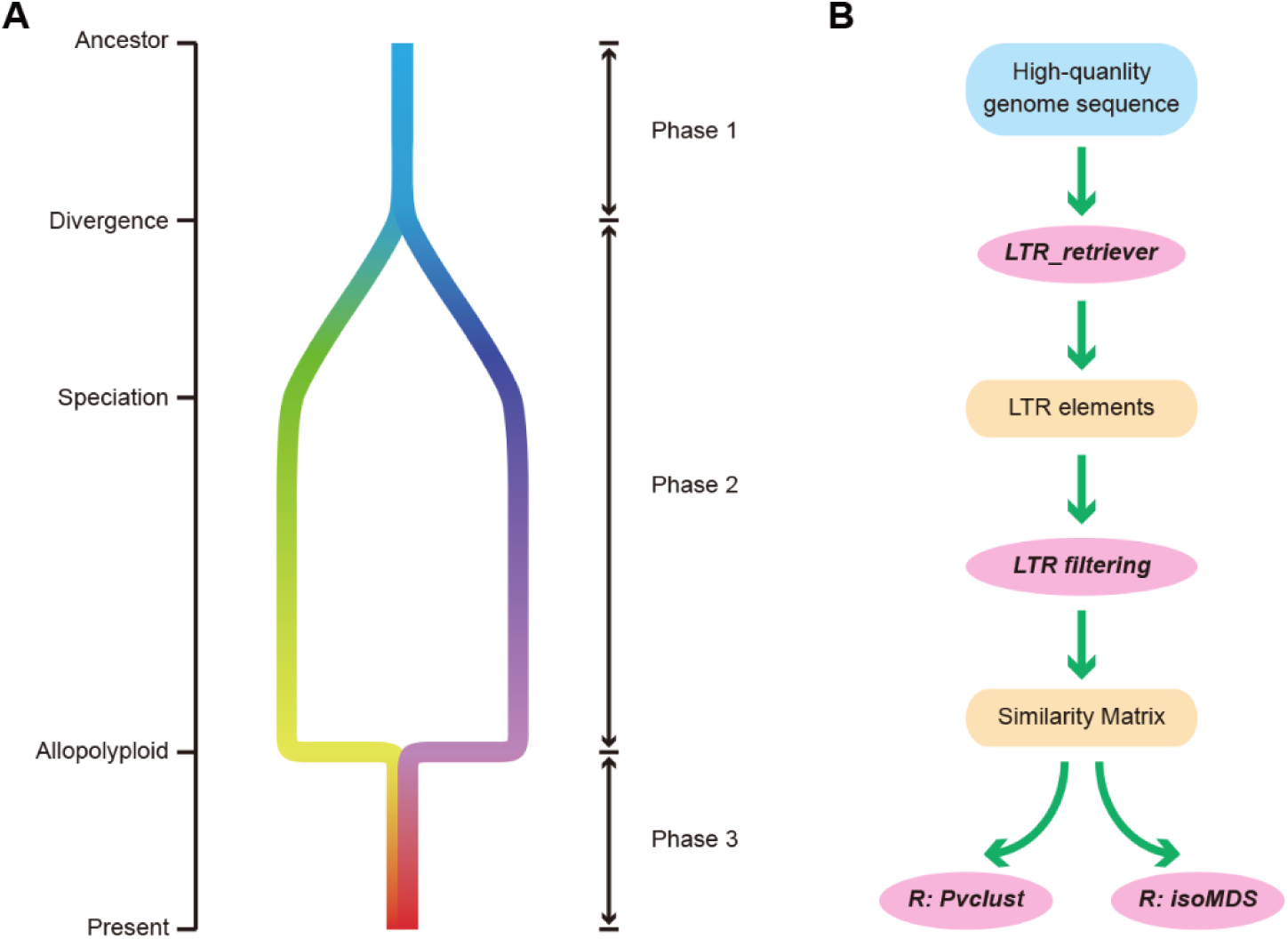
Schematic overview of the Serial Similarity Matrix (SSM) method. (A) Genetic principle: LTR differentiation during the evolutionary process from diploid ancestor genomes to allopolyploid genomes, highlighting the three key phases. (B) Computational pipeline: Genome-wide long terminal repeats (LTRs) identification, similarity matrix construction, subgenome inference using R-based clustering and visualization.

To validate this approach, we selected two well-studied allopolyploid species with known subgenomes: *Eragrostis tef* (AABB) (VanBuren et al. 2020) and *Gossypium hirsutum* (AADD_1_) (Hu et al. 2019) (Supplemental Table S1). Additionally, we applied the SSM method to an artificially created allopolyploid genome, “GmaGso,” which combines the diploid genomes of *Glycine max* and *G. soja* (Xie et al. 2019; Schmutz et al. 2010) (Supplemental Table S1). Phase 2 of *E. tef* was inferred to have occurred 1.1 to 5.0 million years ago (Mya) (VanBuren et al. 2020), while the two progenitors of *G. hirsutum* diverged 6.2 to 7.1 Mya and merged around 1.7 to 1.9 Mya (Hu et al. 2019) (Supplemental Table S1). Following the pipeline outlined in Fig. 1B, we identified and visualized the chromosome clustering within these two different evolutionary periods. The first LTR similarity interval, “90 ∼ 95%”, corresponding to ages of “1.9 ∼ 3.8 Mya”, aligns with Phase 2 (Fig. 1A). The second interval, “98 ∼ 100%”, demonstrates LTR proliferation among chromosomes after allopolyploidization, representing Phase 3 around 0 ∼ 0.8 Mya (Fig. 1A).

The SSM results of Phase 2 displayed complete clustering of chromosomes into the correct subgenomes for both *E. tef* and *G. hirsutum* (Fig. 2A, C), consistent with previous findings (Hu et al. 2019; VanBuren et al. 2020). However, the results of Phase 3 exhibited different patterns between *E. tef* and *G. hirsutum* (Fig. 2B, D). Particularly, the subgenomes of *E. tef* showed less significant clustering compared to those of the cotton genome (Fig. 2B, D). This discrepancy might stem from the lower activity of LTR-RTs in *E. tef* compared to the cotton genome (Supplemental Table S2), indicating a restrained proliferation of LTR-RTs within the subgenome after allopolyploidization. However, it is evident that the significance of clustering in Phase 3 is reduced compared to Phase 2, especially in genomes with low proportions of LTR-RTs (Fig. 2A-D). Thus, using a series of LTR similarity intervals enables inference of the age of allopolyploidy by observing the significance of clustering.

**Figure 2.**
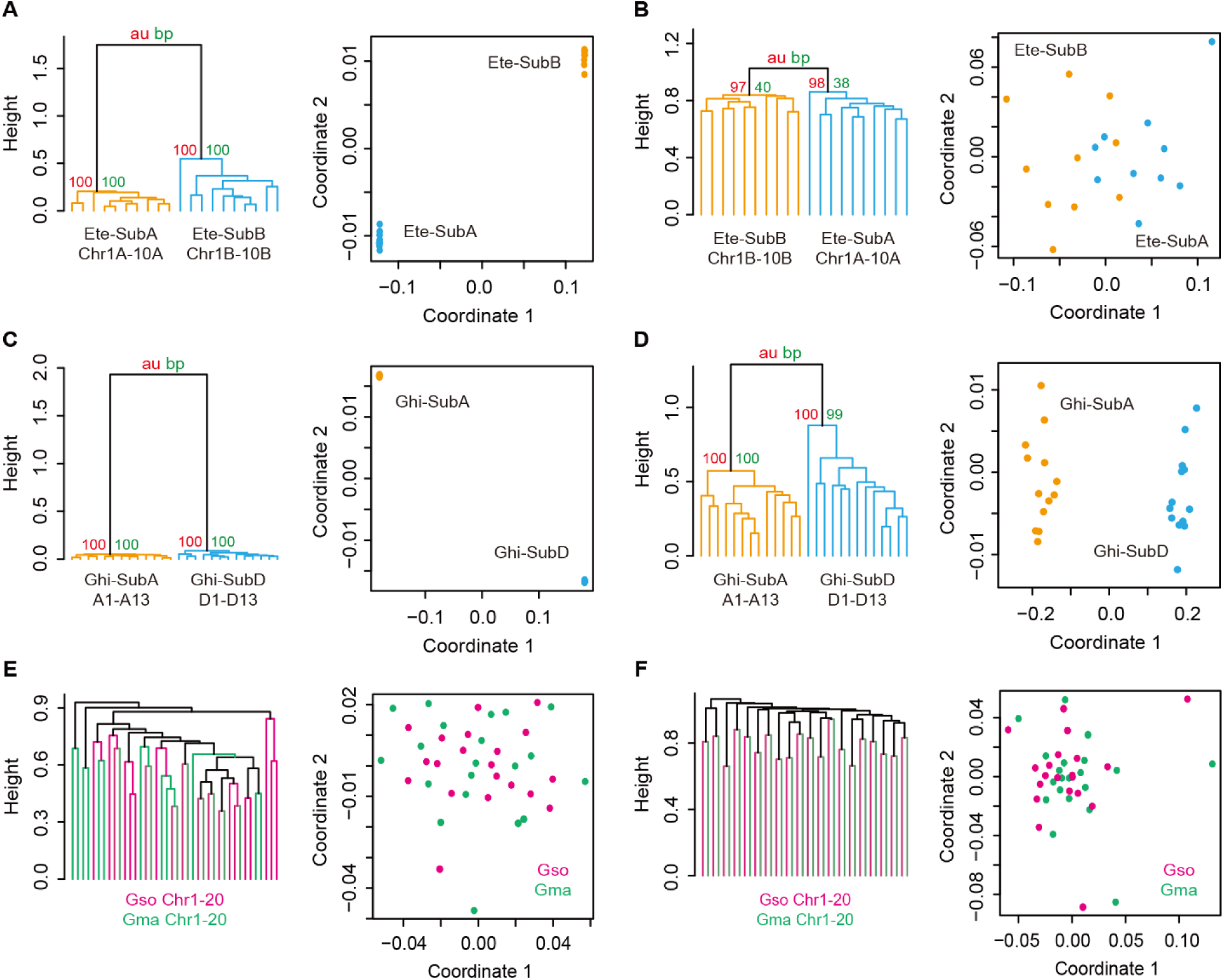
Performances of the SSM method for well-studied allotetraploid genomes and an artificially created allotetraploid genome. (A) SSM clustering results for *Eragrostis tef* with an LTR divergence age of 1.9 - 3.8 Mya. The left panel shows a distance tree generated by the *Pvclust* function, while the right panel displays multidimensional scaling (MDS) clustering. The height of the “*Pvclust*” tree indicates the distance between two samples or clusters. Red and green numbers indicate cluster reliability based on *au* (Approximately Unbiased) and *bp* (Bootstrap Probability) values, respectively. (B) SSM results for *E. tef* with an LTR divergence age of 0 - 0.8 Mya. Note that the two clusters are not significant. (C) SSM results for *Gossypium hirsutum* with an LTR divergence age of 1.9 - 3.8 Mya. (D) SSM results for *G. hirsutum* with an LTR divergence age of 0 - 0.8 Mya. (E) SSM results for GmaGso (*Glycine max* and *Glycine soja*) with an LTR divergence age of 1.9 - 3.8 Mya. No *au* and *bp* values are shown in this case. (F) SSM results for GmaGso (*G. max* and *G. soja*) with an LTR divergence age of 0 - 0.8 Mya.

The divergence of *G. max* and *G. soja* occurred approximately 0.8 Mya (Sedivy et al. 2017; Kim et al. 2010). The artificially created allopolyploid genome GmaGso is expected to exhibit characteristics of Phase 2 as it has not experienced post-allopolyploidization (Phase 3) (Fig. 1A; Supplemental Table S1). SSM analyses identified Phase 1 and Phase 2 in GmaGso (Fig. 1A; Fig. 2E, 2F). In Phase 1, chromosomes from *G. max* and *G. soja* clustered together (Fig. 2E). In Phase 2, the SSM result showed strict pairwise clustering of homoeologous chromosomes between the two *Glycine* species (Fig. 2F). Due to the absence of Phase 3 in the artificial allopolyploid genome GmaGso, subgenome chromosomes did not cluster together. These findings indicate that our SSM method can effectively capture genomic signals of allopolyploidization timing through the distinctive clustering patterns of subgenome chromosomes between pre-allopolyploidization (Phase 2) and post-allopolyploidization (Phase 3).

### SSM analyses for the *Fragaria* genomes

We utilized the SSM method to explore the subgenome partition of the octoploid strawberry genome. Initially, we focused on LTRs that proliferated within the 1.9 ∼ 3.8 Mya interval, corresponding to LTR similarities of 90 to 95%, and constructed a Similarity Matrix (SM) of LTRs across all 28 chromosomes. The resulting clustering, based on SM correlation, identified four distinct subgenomes within the *F.* × *ananassa* genome (Fig. 3A). Notably, while the subgenomes of Fan-Fve and Fan-Fii were consistent with previously findings, the remaining two subgenomes differed from the results of a previous study (Edger et al. 2019) (Fig. 3A).

**Figure 3.**
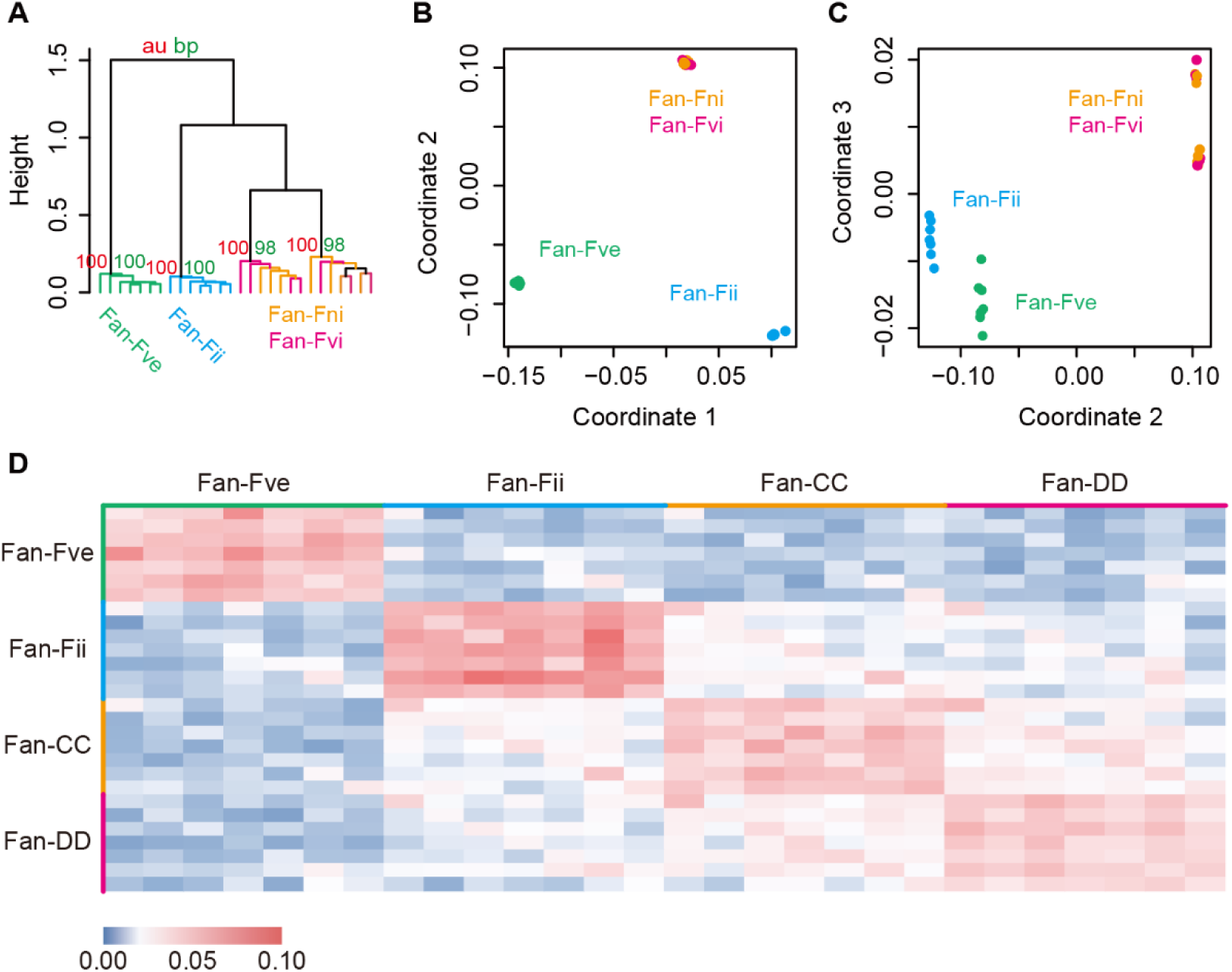
Serial Similarity Matrix (SSM) clustering results of the octoploid strawberry genome. (A) Hierarchical clustering of similarity correlations using *Pvclust*. The similarity boundary was set at 90 - 95%. Subgenomes were highlighted with different colors based on the result of Edger et al. (2019). (B) Multidimensional scaling (MDS) plot based on coordinates 1 and 2. (C) Multidimensional scaling (MDS) plot based on coordinates 2 and 3. (D) Heatmap illustrating the similarity matrix of long terminal repeat (LTR) retrotransposons across the 28 chromosomes.

Multidimensional scaling (MDS) clustering in coordinates 1 and 2 supported three clusters of chromosomes: Fan-Fve, Fan-Fii, and a mixed group of the other two subgenomes (Fig. 3B). MDS clustering in coordinates 2 and 3 separates all four subgenomes, consistent with the correlation clustering results (Fig. 3A, 3C). The SM analysis among 28 chromosomes revealed four clustered subgenome modules, indicating higher similarities within intra-subgenomes than inter-subgenomes (Fig. 3D). Our analysis confirmed the presence of four subgenomes in the octoploid strawberry genome, which we have designated as Fan-Fve, Fan-Fii, Fan-CC, and Fan-DD for subsequent analysis.

We further examined the relationships among five diploid *Fragaria* species and the octoploid strawberry using three intervals of LTR similarities. In the 0 to 1.5 Mya interval, all four subgenomes of *F. × ananassa* clustered together, while the diploid species formed five distinct clusters (Fig. 4A). This analysis revealed the phylogenetic relationships among the diploid species: *F. vesca* ssp. *vesca* and *F. viridis* shared a recent common ancestor, *F. nubicola* and *F. nilgerrensis* were closely related, and *F. iinumae* was positioned as the outgroup of the other four *Fragaria* species (Fig. 4A), consistent with previous phylogenetic studies (Feng et al. 2020; Qiao et al. 2021). In the 1.9 to 3.8 Mya interval, the Fan-Fve subgenome clustered with *F. vesca* ssp. *vesca* and *F. viridis*, while the Fan-Fii subgenome showed a closer relationship with *F. iinumae* than with other diploid species (Fig. 4B). In the 4.2 to 5.8 Mya interval, SSM clustering showed that *F. nubicola* and *F. nilgerrensis* merged into clusters with Fan-Fve, *F. vesca* ssp. *Vesca*, and *F. viridis*, and none of the diploid species in our study emerged as the direct progenitors of Fan-CC and Fan-DD (Fig. 4C). Therefore, our SSM clustering result does not support *F. viridis* as a progenitor species for either Fan-CC or Fan-DD as proposed by Edger et al. (2019). Furthermore, unlike Fan-Fve and *F. vesca* ssp. *vesca*, our analyses did not trace a common ancestor for Fan-Fii and *F. iinumae* (Phase 1). This observation suggests that a subspecies or a close relative of *F. iinumae* may be the direct diploid ancestry rather than *F. iinumae* itself, aligning with a recent finding by Session and Rokhsar (2023). In summary, our investigation revisited the origin of the octoploid genome, and demonstrated the efficacy of the SSM method in unraveling the complex history of allopolyploids in the absence of progenitor information.

**Figure 4.**
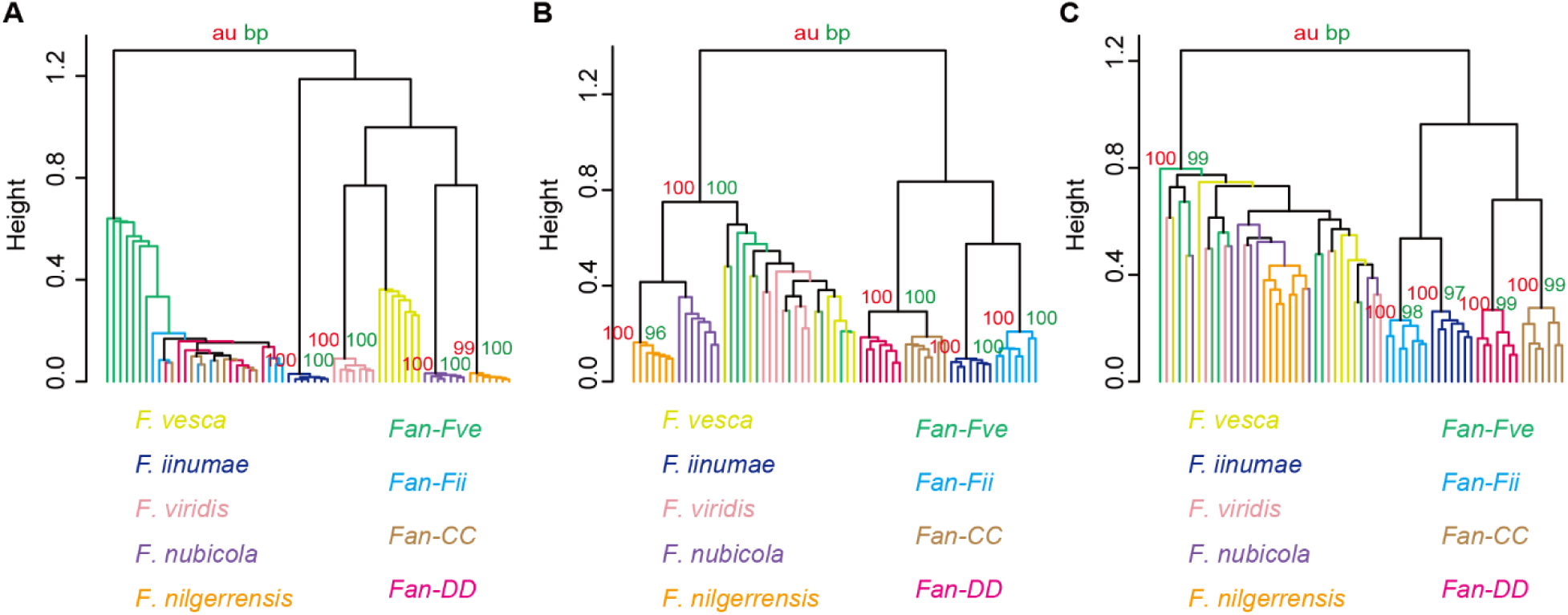
SSM clustering for six *Fragaria species*. Nine genomes or subgenomes representing six *Fragaria* species include *F. vesca* ssp. *vesca, F. iinumae, F. viridis, F. nubicola, F. nilgerrensis*, and the octoploid strawberry (*F.* × *ananassa*) were subjected to SSM clustering analysis within three intervals of LTR age: (A) 0 - 1.5 Mya, (B) 1.9 - 3.8 Mya, (C) 4.2 - 5.8 Mya.

### Identification of diploid progenitors of the octoploid strawberry based on resequencing data

Genomic short reads from five *Fragaria* species (*F. vesca* ssp. *vesca*, *F. vesca* ssp. *bracteata*, *F. iinumae*, *F. nipponica*, and *F. viridis*) (Supplemental Table S3) were aligned to six reference genomes, including *F. vesca* ssp. *vesca* (Shulaev et al. 2011), *F. iinumae* (Edger et al. 2020), and newly assembled subgenomes (Fan-Fve, Fan-Fii, Fan-CC, and Fan-DD), to identify the diploid progenitors of the octoploid strawberry. Alignment coverages revealed that the Fan-Fve subgenome shared greater similarity with *F. vesca* ssp. *vesca* (approximately 80% coverage on average) than with *F. vesca* ssp. *bracteata* (approximately 32% coverage on average) (Supplemental Fig. S1). This contradicts previous inferences based on phylogenetic trees of chloroplast genomes (Njuguna et al. 2013), suggesting a more complex evolutionary history for the Fan-Fve subgenome. Both alignment coverage and genetic distance supported a close relationship between *F. iinumae* and the Fan-Fii subgenome (Supplemental Fig. S1, S2). A relatively low coverage (∼37%) of *F. iinumae* reads could be mapped on its own reference genome, suggesting *F. iinumae* might have undergone substantial intraspecific genome diversification. This observation is consistent with the results obtained from SSM clustering (Fig. 4). In contrast, the Fan-CC and Fan-DD subgenomes showed limited similarities to *F. nipponica* or *F. viridis* (Supplemental Fig. S1, S2). Overall, the progenitor species identified by SSM clustering were largely supported by genomic divergence estimates based on genome sequence comparison between resequencing data and reference genomes.

### Phylogenetic discordance of *Fragaria* species

We examined the phylogenetic relationships among *Fragaria* species by analyzing orthologous genes across five *Fragaria* genomes: *F. vesca* ssp. *vesca*, Fan-Fve, Fan-Fii, Fan-CC, and Fan-DD. A total of 12,046 syntenically conserved gene pairs were identified. Unrooted phylogenetic trees were constructed for these gene pairs, revealing the overall topology relationships among the *Fragaria* species. The consensus tree of these gene trees suggested a close relationship between Fan-CC and Fan-DD (Supplemental Fig. S3).

Analysis of phylogenetic node topologies (Fig. 5A) revealed substantial diversification among the five genomes. Only 33.8% of gene trees supported the sister species relationship of *F. vesca* ssp. *vesca* and Fan-Fve (topology I; Fig. 5A). Approximately 16.3% of gene trees supported a closer relationship between *F. vesca* ssp. *vesca* or Fan-Fve with the other three subgenomes (Fan-Fii, Fan-CC, Fan-DD) (topology II, III, IV, V, VI, VII; Fig. 5A). Notably, a subset of gene trees (topology VIII, IX, X; Fig. 5A) that contain significantly higher percentage of orthologous genes (approximately 22.9% on average) supported the closest relationships among the three subgenomes of Fan-Fii, Fan-CC, and Fan-DD (Fisher’s exact test, P value < 0.05). This pervasive phylogenetic discordance among the three subgenomes suggests that individual gene trees may not accurately reflect their true evolutionary history. Furthermore, the distribution of orthologous genes supporting the discordant topologies VIII, IX, and X across the *F. vesca* ssp. *vesca* reference genome showed no enrichment in specific genomic regions (Fig. 5B). This observation suggests that Fan-Fii, Fan-CC, and Fan-DD might have experienced incomplete lineage sorting or ancestral homoeologous gene exchanges after allopolyploidization events.

**Figure 5.**
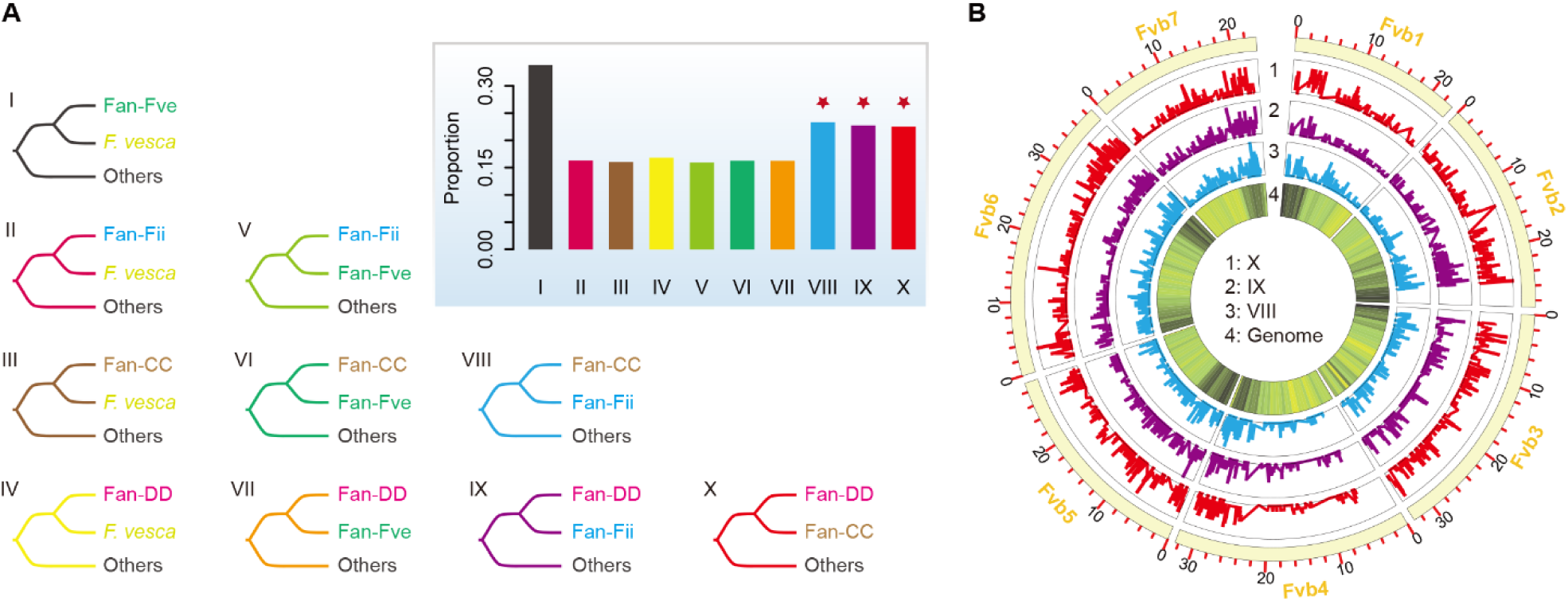
Phylogenetic discordance of syntenic orthologous genes. (A) Phylogenetic topologies of five *Fragaria* genomes. The five genomes include *F. vesca* ssp. *vesca* and the four subgenomes of octoploid strawberry (*Fragaria* × *ananassa*): Fan-Fve, Fan-Fii, Fan-CC, and Fan-DD. The number of topologies with different nodes is presented at the top right. The three topologies marked with red stars (VIII, IX, and X) are significantly enriched compared with the other six (II, III, IV, V, VI, and VII). (B) Genomic distribution of genes exhibiting topologies VIII, IX, and X. These three topologies correspond to those highlighted in panel (A). The density of all syntenic orthologous genes (Track 4) is shown along the genome of *F. vesca* ssp. *vesca*.

### New model for the origin of octoploid strawberry

Syntenic analysis revealed a conserved collinearity pattern between *F. vesca* ssp. *vesca* and the four subgenomes of the octoploid strawberry (Supplemental Fig. S4). While the *Ks* peak (∼0.010) of syntenic genes between *F. vesca* ssp. *vesca* and Fan-Fve subgenome (Supplemental Fig. S5) aligns with the estimated allopolyploidization event timing (∼1.1 Mya) (Njuguna et al. 2013), resolving relationships among other subgenomes using *Ks* alone proved difficult (Supplemental Fig. S5).

To better understand the evolutionary history among genomes, we estimated the lower limit of “Phase 2” LTR ages by assessing sequence divergence of the best-hit LTRs, calibrated by the neutral mutation rate of LTRs. Subsequently, we employed SSM analyses to disentangle the evolutionary history among these genomes as well as the complex allopolyploidization history of the octoploid strawberry. SSM analysis of six genomes (*F. vesca*, *F. iinumae*, Fan-Fve, Fan-Fii, Fan-CC, and Fan-DD) across five time intervals (Fig. 6) revealed distinct evolutionary stages:

**4.2 - 5.8 Mya:** The progenitors of Fan-CC and Fan-DD subgenomes diverged into two independent diploid species. Their common ancestor dates back more than 5.8 million years ago (Fig. 6E, F). The common ancestor of all six genomes likely predates this period.

**3.1 - 4.2 Mya:** SSM analysis completely separated Fan-CC and Fan-DD subgenomes in this time frame (Fig. 6D), suggesting the first polyploidization event of the octoploid strawberry occurred during this time frame.

**1.9 - 3.1 Mya:** *F. iinumae* and Fan-Fii shared similar LTRs in this time frame (Fig. 6C), supporting a polyploidization event involving *F. iinumae* and the ancestral allotetraploid during this interval.

**0.8 - 1.9 Mya:** Fan-Fve, and *F. vesca* genomes diverged from the other three subgenomes (Fan-Fii, Fan-CC, and Fan-DD) (Fig. 6B), suggesting that a polyploidization event between *F. vesca* and the ancestral hexaploid occurred in this interval, consistent with the previous estimation of ∼1.1 Mya (Njuguna et al. 2013).

**0 - 0.8 Mya:** The *F. iinumae* genome remained distinct from the four subgenomes of strawberry (Fig. 6A). The Fan-Fve subgenome began accumulating LTRs similar to the other three subgenomes (Fan-Fii, Fan-CC, and Fan-DD), diverging from its *F. vesca* progenitor (Fig. 6A).

**Figure 6.**
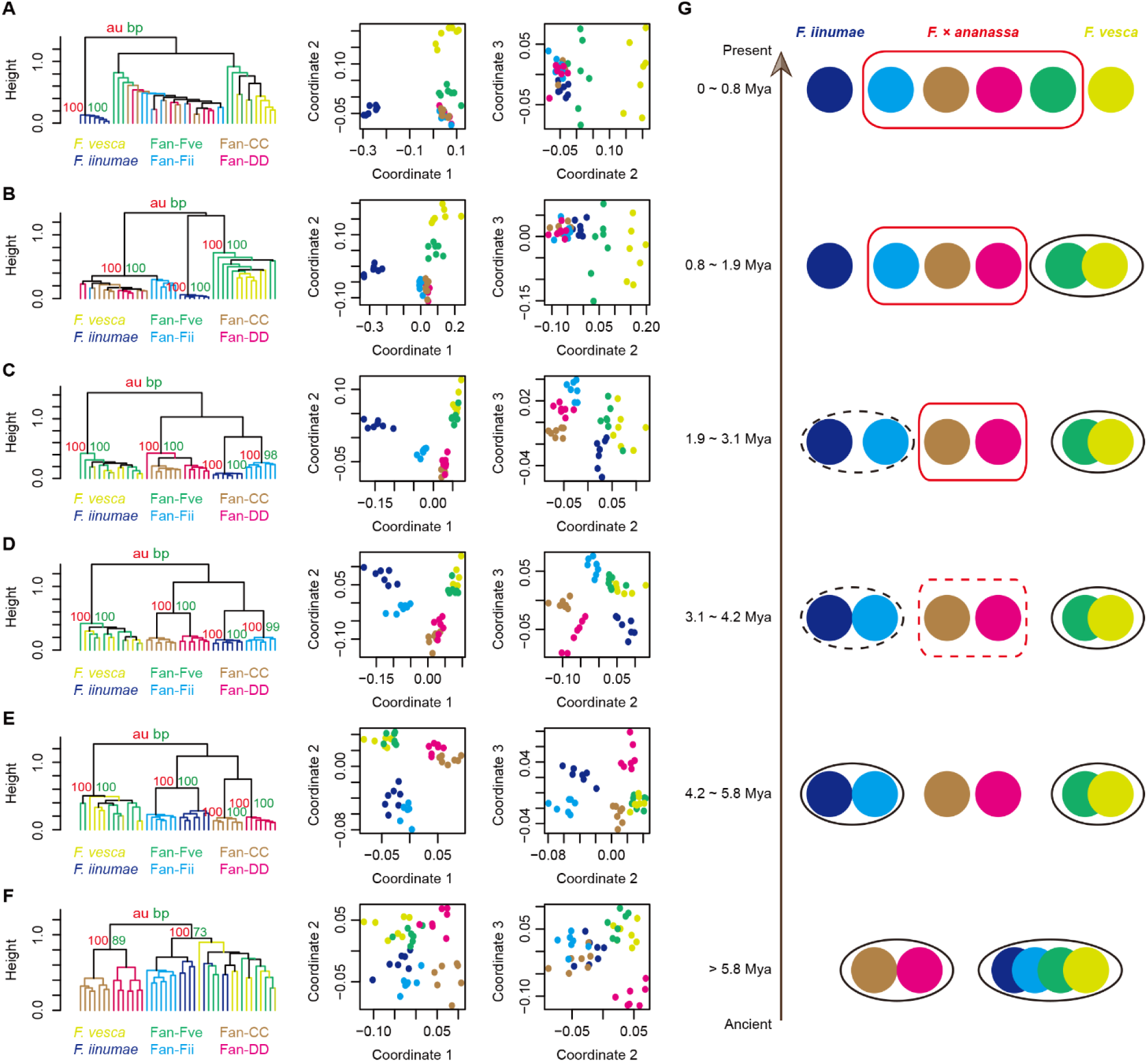
SSM analyses and evolutionary model of *Fragaria* species. (A) SSM clustering between 0 - 0.8 Mya. Two diploid *Fragaria* species (*F. vesca* ssp*. vesca* and *F. iinumae*) and the octoploid strawberry (*F. × ananassa*) were analyzed. The lower boundary of LTR ages were estimated from the sequence divergence between best-hit LTRs and the neutral mutation rate of LTRs. The top hierarchical cluster with higher *au* and *pb* values than all the bottom clusters were retained for modeling analyses. The two MDS plots show the clustering of coordinate 1 vs. coordinate 2 and coordinate 2 vs. coordinate 3. (B) SSM clustering between 0.8 - 1.9 Mya. (C) SSM clustering between 1.9 - 3.1 Mya. (D) SSM clustering between 3.1 - 4.2 Mya. (E) SSM clustering between 4.2 - 5.8 Mya. (F) SSM is clustering more than 5.8 Mya. (G) A new scenario for the origin of octoploid strawberry. Each solid circle presents one genome or subgenome. Genomes within the same frame share a common ancestor. The dashed frame indicates that the genomes are not completely diverged. The red frame highlights the different evolutionary stages of octoploid strawberry, while the black frame shows the relationship between progenitor species and strawberry subgenomes.

Our findings lead us to propose a new model for the evolutionary history of the octoploid strawberry (Fig. 6G). This model suggests three key polyploidization events: firstly, the most ancient event occurred between the progenitor species of Fan-CC and Fan-DD around 3.1 to 4.2 Mya; secondly, the origin of the hexaploid ancestor containing Fan-Fii, Fan-CC, and Fan-DD subgenomes took place approximately 1.9 to 3.1 Mya; and finally, the merging of the progenitor of *F. vesca* with the ancestral hexaploid genome occurred around 0.8 to 1.9 Mya (Fig. 6G).

Furthermore, our analyses shed light on the phylogenetic relationships among the six genomes. Fan-CC and Fan-DD shared a relatively recent common ancestor, while Fan-Fve and Fan-Fii had an even more recent common ancestor (Fig. 6G), supported by an unrooted consensus tree of the gene trees of their syntenic homologs (Supplemental Fig. S3), which is consistent with previous studies (Qiao et al., 2021).

## Discussion

The exploration of genome evolutionary dynamics is pivotal for understanding the genetic mechanisms behind adaptation, speciation, and biodiversity. Whole genome duplication (WGD) events have played a significant role in the evolution of many organisms, including plants and animals. Nearly all seed plants have experienced at least one ancient WGD event (Tang et al. 2008; Jaillon et al. 2007; Bowers et al. 2003; Vanneste et al. 2014). Subsequent differential retention and divergence of duplicated genes have played an essential role in shaping genetic diversity. Therefore, accurate subgenome partition is essential for deciphering genome evolution and adaptation. Accurate subgenome partition enables the reconstruction of ancestral genomes, shedding light on the genomic changes that have occurred over evolutionary time. This information not only enhances our understanding of evolutionary processes but also provides valuable insights into the genetic basis of phenotypic diversity and adaptation. Moreover, accurate subgenome partition is essential for unraveling the genetic architecture underlying complex traits and facilitating comparative genomics analyses across diverse species.

Advancements in sequencing technologies have facilitated comprehensive evolutionary genomics studies. However, accurate subgenome partition remains challenging, particularly when natural diploid progenitors cannot be identified. In this study, we introduced a novel approach utilizing long terminal repeat retrotransposons (LTR-RTs) coupled with the Serial Similarity Matrix (SSM) method to elucidate the evolution of allopolyploid genomes. LTR-RTs, constituting the most abundant group of transposable elements in plant genomes, offer a potent tool for elucidating the evolutionary history of speciation owing to their propensity for “jumping within the genome”. This unique characteristic enables the detection of sequence similarities between chromosomes within subgenomes at specific evolutionary time frame, facilitating subgenome clustering.

The accuracy and reliability of our SSM method were confirmed through its application to well-studied alloploidy genomes and an artificial alloploidy genome. The results align with previous studies, further validating the effectiveness of our SSM method. Moreover, the SSM method provides compelling visualizations that reveal the evolutionary processes of allopolyploidization. By estimating the lower limits of LTR age through sequence divergence between best-hit LTRs, the SSM method offers valuable insights into the timing of allopolyploidization events.

We applied the SSM method to investigate the octoploid strawberry (*Fragaria × ananassa* Duchesne) genome, a species whose evolutionary origin has long been a subject of debate among researchers. The octoploid strawberry presents a particularly challenging case due to the lack of knowledge regarding its progenitors. Phylogenetic relationships of *Fragaria* species were inferred using nuclear and plastid markers (Dillenberger et al. 2018; Composition et al. 2017; Njuguna et al. 2013; Qiao et al. 2021; Rousseau-Gueutin et al. 2009). Edger *et al*. (Edger et al. 2019) reported the four subgenomes of the octoploid strawberry and inferred their progenitor species based on gene trees. However, the reliability of these findings was called into question due to methodological limitations and incomplete knowledge of progenitor species (Feng et al. 2020; Liston et al. 2020). In our study, revisiting subgenome clustering using the SSM method revealed widespread discordances in phylogenetic relationships among *Fragaria* species, likely stemming from incomplete lineage sorting and homologous exchanges among subgenomes. Importantly, our analysis identified *F. vesca* and *F. iinumae* as closely related species for two of the subgenomes, while challenging the inclusion of *F. nipponica* and *F. viridis* as progenitors of the octoploid strawberry. Furthermore, our analysis suggests that *F. vesca* ssp. *vesca* is genomically closer to the Fan-Fve subgenome, contradicting previous chloroplast-based models suggesting a potential contribution of *F. vesca* ssp. *bracteata* to one subgenome of the domesticated strawberry (Njuguna et al. 2013). Similarly, our analysis suggests that *F. iinumae* and the subgenome of Fan-Fii are closely related but couldn’t trace back to the common ancestor of Fan-Fii and *F. iinumae*, suggesting the possibility of an extinct *F. iinumae*-like diploid species as the direct donor of Fan-Fii, consistent with earlier hypotheses based on chromosomal structure (Liu et al. 2016; Wei et al. 2017).

In addition, our analysis inferred three allopolyploidization events (0.8 - 1.9 Mya, 1.9 - 3.1 Mya, and 3.1 - 4.2 Mya) contributing to the octoploid strawberry genome. The progenitors of Fan-CC and Fan-DD merged first, followed by the most recent addition of Fan-Fve. This revised understanding of the evolutionary history of octoploid strawberry provides essential insights into the complex allopolyploidization events that shaped this species.

## Methods

### Genetic basis and pipeline design of the SSM method

Figure 1 illustrates the genetic basis and computational pipeline of the SSM method. We conceptualize genomic differentiation of LTR retrotransposons in three phases. Phase 1 represents the stage before the progenitor species diverge from each other. During the phase 2, divergence and speciation of progenitor species occurred (Seehausen et al. 2014; Feder et al. 2012; Wu and Ting 2004) (Fig. 1A). LTR retrotransposon diversification should coincide with speciation events. During phase 3, allopolyploidization occurred and the two progenitor genomes merged in a polyploid nucleus, leading to shared transposable element activities (Steige and Slotte 2016; Parisod et al. 2010; Edger et al. 2018) (Fig. 1A). Based on this scenario, subgenome-specific signatures are identifiable during Phase 2 (Fig. 1A). LTR retrotransposons proliferating in this phase are expected to be subgenome-specific (Fig. 1A). We estimated the lower limit of insertion age for each incomplete LTR using the neutral mutation rate for LTRs (μ = 1.3 × 10^-8^ site^-1^·year^-1^) and sequence divergence between best-hit LTRs. For allopolyploid genomes, the lower limits of LTR ages are used to infer the lower bound for Phase 2 and thus, the allopolyploidization event.

The genome-wide identification of LTRs was performed using LTR_retriever, LTR elements, and LTR filtering (Fig. 1B). The filtered LTRs were then subjected to the SSM analysis. The similarity matrix (SM) of LTRs were calculated as below:

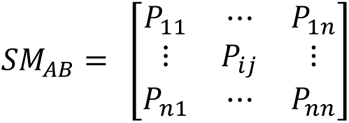

The SM_AB_ presents a similarity matrix of an allopolyploid genome (AABB); n means the total number of chromosomes of both A and B subgenomes; and P_ij_ is the best-hit alignments of LTR pairs between i and j chromosomes divided by the actual numbers of LTRs on i and j. This similarity matrix (SM_AB_) was subsequently used in clustering analyses using either correlation or Euclidean distance of SM.

### Validating the SSM method using well-studied allopolyploid genomes

We applied the SSM method to assess subgenome identification in well-studied allotetraploid genomes: *Eragrostis tef* (AABB) (VanBuren et al. 2020) and *Gossypium hirsutum* (AADD1) (Hu et al. 2019). To further validate the method, we created an artificial allotetraploid genome (“GmaGso”) by merging the genomes of two closely related diploid species, *Glycine max* and *G. soja* (Xie et al. 2019; Schmutz et al. 2010), and applied the SSM method to assess subgenome identification in this artificial allotetraploid genome. Chromosome-level genome sequences for these species were obtained from Cottongen (Yu et al. 2014), SoyBase (Grant et al. 2009), and NCBI.

Subsequently, LTR retrotransposons (LTR-RTs) in each genome were identified using the *LTR_retriever* pipeline (Ou and Jiang 2018). The raw dataset of full-length LTR-RTs was identified using both *LTRharvest* and *LTR Finder*, and then filtered by *LTR_retriever* (Ellinghaus et al. 2008; Xu and Wang 2007). *LTRharvest* was executed with the parameters “-similar 80 -motif tgca -motifmis 1 -minlenltr 100 -maxlenltr 7000 -mintsd 2 -maxtsd 6”, and *LTR Finder* with “-D 15000 -d 1000 -L 7000 -l 100 -p 20 -C -M 0.80”. The resulting set of LTR elements was then subjected to similarity searches using BLAST.

All-against-all LTR comparisons were performed in the three genomes (two natural and one artificial allopolyploid). The comparisons were filtered based on the following criteria: 1) the length of LTR elements >= 100 bp; 2) E-value <= 1E-10; 3) only retaining best-hits of all-against-all comparisons for sequence similarity matrix (SM) construction. The number of best-hit LTR pairs within the allopolyploid genomes was recorded for specified intervals of LTR similarity. LTR similarity intervals for constructing the SM were chosen to encompass Phase 2 divergence (Fig. 1A). This is distinct from previous methods (Jia et al. 2022; Session and Rokhsar 2023; Wang et al. 2022). The serial intervals for an unknown allopolyploid genome could be set with overlapping ranges with a recommended interval size of approximately 2-4% similarity. The interval size can be adjusted based on the genome-wide LTR-RT content and clustering patterns. For *G. hirsutum*, *E. tef*, and GmaGso, these intervals were established as 90-95% and 98-100% similarity.

Within each interval, SMs were generated by counting best-hit LTRs between chromosome pairs, calibrated by the total number of LTR in each chromosome pair. The resulting SMs were then subjected to clustering and visualization using R-based methods, “*Pvclust*” (Suzuki and Shimodaira 2006) and “*isoMDS*” (Venables and Ripley 2002). “*Pvclust*” clustered chromosomes based on correlations, while “*isoMDS*” employed Euclidean distance for clustering chromosomes into subgenomes. Reliability of chromosome clusters could be assessed using the approximately unbiased (au) and bootstrap probability (bp) values derived from 1000 bootstraps by “*Pvclust*”.

### Performance of SSM in the octoploid strawberry genome

The octoploid strawberry genome (*Fragaria* × *ananassa*) has been sequenced and analyzed (Edger et al. 2019), but identification of its subgenomes due to uncertain progenitor species has been a topic of ongoing revision (Liston et al. 2020; Session and Rokhsar 2023). In this study, we reassessed the subgenomes of the octoploid strawberry using the SSM approach, following the pipeline described above. The SM was constructed with a similarity boundary set at “90 - 95%”. After clustering and visualizing chromosome similarities, we compared our subgenome definitions with those previously reported (Edger et al. 2019; Session and Rokhsar 2023).

We applied the SSM method to investigate the possible progenitor species contributing to the four subgenomes of octoploid strawberry. Five diploid *Fragaria* species - *F. vesca* ssp. *vesca*, *F. iinumae*, *F. viridis*, *F. nubicola*, and *F. nilgerrensis vesca* (Shulaev et al. 2011; Edger et al. 2020; Feng et al. 2020) - were included in the SSM analysis alongside the octoploid strawberry (*F. × ananassa*). We generated SSMs using three similarity intervals (96-100%, 90-95%, and 85-89%) to correspond with estimated divergence times (0-1.5 Mya, 1.9-3.8 Mya, and 4.2-5.8 Mya, respectively). Clustering analyses were then performed across these six species, encompassing nine genomes or subgenomes in total.

### Progenitor inference using the raw reads of five *Fragaria* species

In a previous study, four diploid progenitors of the octoploid strawberry (*Fragaria* × *ananassa*) were proposed: *F. vesca* ssp. *bracteata*, *F. iinumae*, *F. nipponica*, and *F. viridis* (Edger et al. 2019). To independently validate these findings, we obtained raw genomic resequencing reads for five diploid *Fragaria* species: *F. vesca* ssp. *vesca*, *F. vesca* ssp. *bracteata*, *F. iinumae*, *F. nipponica*, and *F. viridis* (Supplemental Table S3). To reassess the proposed progenitors, we mapped the raw reads to six reference genomes: *F. vesca* ssp. *vesca* (Shulaev et al. 2011), *F. iinumae* (Edger et al. 2020), and the four subgenomes of the octoploid strawberry (Fan-Fve, Fan-Fii, Fan-CC, Fan-DD) derived from our SSM subgenome separation method. The *F.* × *ananassa*, *F. vesca* ssp. *vesca* and *F. iinumae* reference genomes were obtained from the Genome Database for Rosaceae (GDR, https://www.rosaceae.org) (Jung et al. 2019). Raw reads were aligned to each reference genome using the BWA aligner (Li and Durbin 2009), followed by processing with SAMTOOLS (Li et al. 2009) and PICARD (http://broadinstitute.github.io/picard). Genotype calling and filtering were performed using the GATK package with default parameters (McKenna et al. 2010).

For each raw read dataset and reference genome pair, we calculated two parameters: 1) Proportion of Reference Covered: The percentage of each reference chromosome covered by aligned reads; and 2) Genetic Distance: The genetic distance between each species pair, estimated from mismatches in the filtered genotypes. Specifically, if two alternative bases were observed at a site (AA), the distance increased by 1; a single alternative base (Aa) led to 0.5 increment of the genetic distance. The total genetic distance was then normalized by the length of the covered alignment region.

### Phylogenomic analysis of syntenic genes among strawberry subgenomes

We explored the phylogenetic relationships of protein-coding genes among the four subgenomes of octoploid strawberry. Syntenic blocks across the diploid *F. vesca* ssp. *vesca* genome and the four subgenomes of the octoploid *F.* × *ananassa* (Fan-Fve, Fan-Fii, Fan-CC, Fan-DD) were identified using PyMCScan (Wang et al. 2012). For each syntenic gene pair, coding sequences (CDS) were aligned using MUSCLE (v3.8.31) (Edgar 2004). Synonymous substitution rates (Ks) were then calculated using KaKs_Calculator 2.0 (Wang et al. 2010).

PyMCScan was used to extract syntenic orthologous genes from each of the five genomes. Identified syntenic orthologs were aligned at the nucleotide level, guided by their corresponding protein alignments generated with MUSCLE (v3.8.31) (Edgar 2004). Maximum Likelihood (ML) phylogenetic trees were reconstructed using FastTree (Price et al. 2010). Phylogenetic discordance across the *F. vesca* ssp. *vesca* genome was assessed by analyzing the resulting ML trees. The relative abundance of different tree topologies was visualized using DensiTree (Bouckaert 2010). The proportions of each topology among the five genomes were quantified, the density of discordant topologies along the *F. vesca* ssp. *vesca* genome was plotted using TBtools (Chen et al. 2020).

### Evolutionary history of the octoploid strawberry genome

Our SSM method elucidates the origin of the octoploid genome and provides age estimates for the three allopolyploidization events. We applied the SSM method to the octoploid *F.* × *ananassa* genome and its two known progenitor species, *F. vesca* ssp. *vesca* and *F. iinumae*. Six intervals of LTR similarity were set for SSM analysis (≤ 85%, 85 - 89%, 89 - 92%, 92 - 95%, 95 - 98%, and 98 - 100%), corresponding to six time periods ranging from present to more than 5.8 million years ago (Mya). The 42 chromosomes of *F.* × *ananassa*, *F. vesca* ssp. *Vesca*, and *F. iinumae* were clustered and visualized for each of the six time periods. This analysis allowed us to model the evolutionary relationships among the six *Fragaria* genomes or subgenomes from the three species (*F.* × *ananassa*, *F. vesca* ssp. *vesca,* and *F. iinumae*).

### Data access

The code used to generate results is available at https://github.com/HaominLyu/SSM.git.

## Competing interest statement

The authors declare no competing interests.

## Acknowledgments

This work was supported by National Institute of Food and Agriculture (NIFA) – Specialty Crop Research Initiative (SCRI) Grant 2022-51181-38241 to QY.

## Author contributions

H.L. designed and performed experiments and wrote the manuscript. S.O. and W.C.Y. revised the manuscript. Q.Y. provided supervision, guidance, and support to H.L. and supervised the project and wrote the manuscript.

## Notes

### Competing Interest Statement

The authors have declared no competing interest.

